# Chemically-inducible CRISPR/Cas9 circuits for ultra-high dynamic range gene perturbation

**DOI:** 10.1101/2024.12.27.630546

**Authors:** Rajini Srinivasan, Tao Sun, Azalia Sandles, Diana Wu, Liang Wang, Amy Heidersbach, Clark Ho, Shiqi Xie, Andrew Ng, Benjamin Haley

**Affiliations:** Department of Molecular Biology, Genentech Inc., South San Francisco, CA; Department of Functional Genomics, Genentech Inc., South San Francisco, CA

**Author notes:** equal contribution (Rajini Srinivasan, Tao Sun). Centre de Recherche de l’Hôpital Maisonneuve-Rosemont, Pavillon Claudine D’Amours, 5690 Boul. Rosemont, Montréal, QC, Canada H1T 2H2.

## Abstract

CRISPR/Cas9 technologies provide unique capabilities for modeling disease and understanding gene-to-phenotype connections. In cultured cells, chemical-mediated control of Cas9 activity can limit off-target effects and enable mechanistic study of essential genes. However, widely-used Tet-On systems often show “leaky” Cas9 expression, leading to unintended edits, as well as weak activity upon induction. Leakiness can be distinctly problematic in the context of Cas9 nuclease activity, which may result in cumulative DNA damage and degradation of the target cell genome over time. To overcome these deficiencies, we established transgenic platforms that minimize Cas9 functionality in the off-state along with maximized and uncompromised on-state gene editing efficiency. By combining conditional destabilization and inhibition of Cas9, we developed an all-in-one (one or multiple guide RNAs and Cas9) ultra-tight, Tet-inducible system with exceptional dynamic range (on vs. off-state) across various cell lines and targets. As an alternative to Tet-mediated induction, we created a branaplam-regulated splice switch module for low-baseline and robust Cas9 activity control. Lastly, for circumstances where DNA damage needs to be avoided, we constructed a dual-control, Tet-inducible CRISPRi module for tight and potent transcriptional silencing. This upgraded suite of inducible CRISPR systems has broad applications for numerous cell types and experimental conditions.

## Introduction

CRISPR/Cas9 is a highly adaptable and efficient genome editing tool. Commonly, users design single guide RNAs (sgRNAs) that direct Cas9 to generate double-stranded DNA (dsDNA) breaks at specified genomic target sites. Inaccurate repair of the Cas9-mediated DNA damage results in a permanent genetic modification, which allows for disruption of individual genes or thousands in the context of functional genomic screens. The power of this technology is substantially increased through methods that enable users to manipulate the timing and duration of Cas9 expression. Such control helps to minimize off-target DNA damage, and facilitates the study of gene function(s) over a wide range of conditions and cellular states. Inducible gene disruption also enables mechanistic and temporal assessment of cell populations depleted of essential targets, which, due to the lethal phenotypes, present challenges with approaches that rely on constitutive Cas9 expression. However, any inducible Cas9 nuclease system is dependent on low or essentially no background editing in the “off” state, since this would result in an accumulation of permanent gene edits over time ^1,2^. In addition, the post-induction “on” state should provide high and uniform targeting efficacy across the desired cell population. Ideally, such a system would function consistently over a wide range of cell types and target loci.

Broadly applied for both *in vitro* and *in vivo* experiments, Tetracycline (Tet)/Doxycycline (Dox)-inducible systems have shown promise for timed activation of Cas9 activity. Yet, the dynamic range (e.g., contrast between the on and off state editing) of Tet-based induction is often limited by issues such as “leaky” background expression or sub-optimal gene targeting efficiency after induction ^3,4^. Clonal selection from transgenic cell populations can help mitigate basal Cas9 activity, but this approach is labor-intensive, risks skewing genetic and phenotypic diversity representative of the parental cells, and may fail to fully address variability across cell types. Alternative strategies for induced gene editing have been explored, including integration of a Tet-On/Cas9 cassette with insulators separating the promoter and the reverse Tet transactivator (rtTA) elements into genomic safe harbor sites ^5^, chemical stabilization/destabilization of degron-linked Cas9^4,6–9^, split-Cas^10^, intein-Cas9^11^, hydroxytamoxifen inducible Cas9^12,13^ and Dox-inducible sgRNA ^14,15^ or combinations of each^4,6,8^. Despite showing value, these approaches struggle with challenges related to incomplete induction and/or residual background activity, highlighting the need for technical improvement to achieve peak precision, control, and potency of genome editing.

To address these limitations, we developed a suite of enhanced, inducible CRISPR/Cas9 systems offering full on-state gene perturbation with near-absent background editing. In addition, we provide vectors that enable streamlined, single-step production of cell populations with optimal Cas9 activity profiles, eliminating the need for single cell cloning. For general applications, we engineered an all-in-one, “ultra-tight” transgene that combines Dox-inducible and degron-destabilized Cas9, a separate degron-linked anti-CRISPR module, and a constitutive sgRNA expression module to achieve ∼60-95% editing efficiency and marginal (often unmeasurable) background across multiple genes and cell lines. We further expanded the use of this system through co-expression of two sgRNAs for induced, population-level excision of a >2 kilobase (kb) genomic segment; facilitating conditional deletion of individual repetitive genomic regions, non-coding loci, and more. For circumstances where Tet/Dox-based induction must be avoided, we created an all-in-one, chemically-controlled anti-CRISPR-integrated splice switch module for Tet/Dox-independent regulation of Cas9 activity. Finally, in recognition of experimental conditions that would benefit from transcriptional silencing (e.g. CRISPR inhibition/CRISPRi) rather than directed DNA damage, we established a Dox-inducible, degron-tagged Zim3-dCas9 expression vector that provides superior gene suppression compared to the commonly used KRAB-(KOX1) dCas9 and minimal background silencing.

Together, these tools offer the highest possible contrast between induced and background Cas9 activity *in vitro*, enabling precise, on-demand CRISPR applications that bolster the flexibility and reliability of gene editing in cell-based experiments.

## Results

### The Tet-On CRISPR/Cas9 system exhibits significant basal editing

To establish a baseline for background versus drug-induced editing with a foundational Tet-On Cas9 variant, we used piggyBac to stably integrate an “all-in-one” transgene across a panel of seven human cell lines. Here, *S. pyogenes* (Sp) Cas9 was placed under the control of the TRE3G Tet-inducible promoter upstream of a U6 promoter for constitutive expression of an sgRNA targeting *CD81.* A separate constitutive promoter (EF1α) is used to express a reverse tetracycline transactivator (rtTA) and puromycin resistance (PuroR) gene, which are separated by a T2A linker. Hereafter we term this construct, inclusive of the *CD81* sgRNA, “pTET”. Disruption of *CD81*, encoding a ubiquitously-expressed cell surface protein, can be easily quantified through antibody stain and flow cytometry (Fig. 1A). After generation of stable polyclonal lines with pTET, cell populations were either treated with Dox to induce *CD81* knockout or left untreated as a control. After ten days with or without induction, cells were stained using anti-CD81 and protein levels were monitored by flow cytometry. A representative gating strategy is shown for HEK293T (Fig. S1A). We observed significant and reproducible Dox-independent editing of *CD81* in all tested cell lines (Fig. 1B). Background targeting levels ranged from a low of ∼4% in K562 (bone marrow) cells to as high as 60% in SW480 (colorectal) cells. Upon treatment with Dox, robust (≥95%) depletion of CD81 was observed in all but one cell line (HCT116, ∼80% depletion). Despite potent target disruption, these data highlight a pervasive problem of leaky Cas9 expression in the context of a conventional Tet-On system.

**Figure 1.**
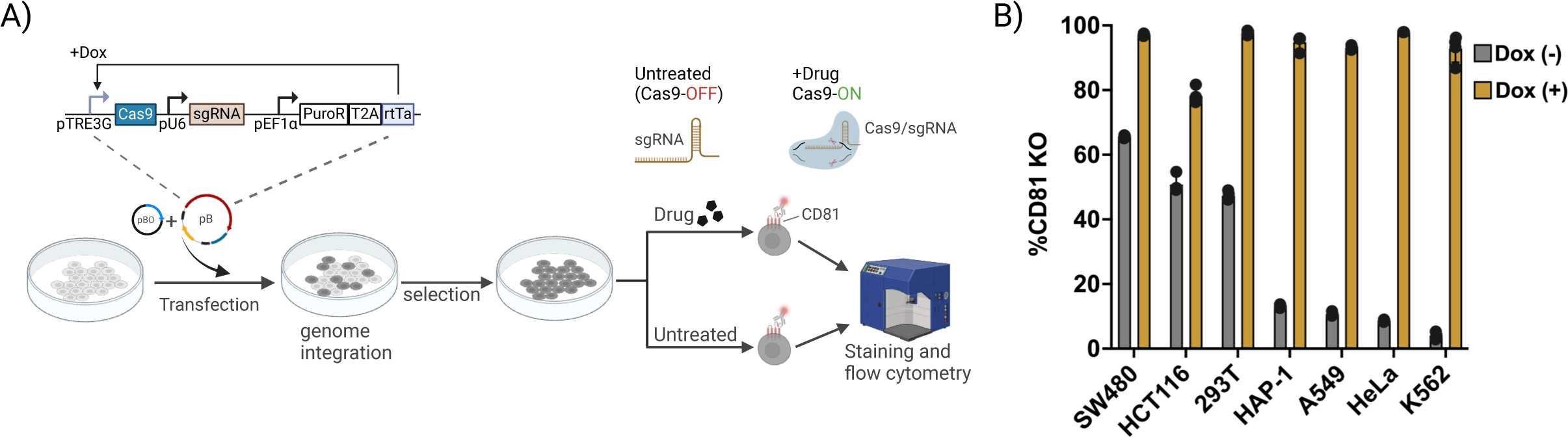
Leaky editing with the pTET system across cell lines. (A) Schematic representation of the pTET piggyBac transposon plasmid (pB) and workflow depicting stable cell line generation with the piggyBac system. Cas9 expression is driven by a TRE3G Tetracycline-inducible promoter (pTRE3G). A human U6 promoter (pU6) drives constitutive expression of the *CD81* sgRNA. Puromycin resistance gene (Puro^R^) and the reverse tetracycline transactivator (rtTA) are constitutively expressed from an EF1α promoter and separated by a self-cleaving T2A linker. Cells that have successfully integrated the cassette are enriched by puromycin selection. Stable lines were treated with Cas9 inducing drug(s) or left untreated as a control, followed by flow cytometry to measure CD81 levels. pBO: PiggyBac transposase. (B) The pTET plasmid was stably integrated into HEK293T, SW480, HeLa, HAP-1, A549, HCT116 and K562 parental cells. Stable cells were treated with 2μg/ml Doxycycline (Dox) for 10 days. Percentage knockout was determined by measuring CD81 negative cells by flow cytometry in untreated and Dox treated cells. Data are shown as mean values +/− SD from at least three independent experiments.

### An ‘ultra-tight” TET-CRISPR/Cas9 system with near-absent basal editing and high knockout efficiency

In order to minimize background editing, we applied a series of modifications to pTET. These adopted different strategies to regulate Cas9 in the uninduced state including conditional stabilization of Cas9 protein, modulated anti-CRISPR expression for directed control of Cas9 activity, and drug-regulated alternative splicing to influence translation of the Cas9-containing pre-mRNA (Fig. 2A-D). The various transgenic circuits, unique control modules, and the specific drug(s) required for Cas9 expression/function are defined in Table 1. For each of these modified genetic circuits, we generated stable polyclonal cell lines in contexts that exhibited high (293T), mid (A549) and low (K562) levels of basal editing. Each stable cell population was treated with the appropriate drug or drug combination for ten days to induce CD81 depletion, while an equal number of cells were left untreated as a control. In order to obtain a performance metric for each circuit that encompasses both ON and OFF-state editing, we determined the dynamic range of Cas9 activity, defined as the ratio between maximum possible knockout efficiency versus leakiness in the uninduced state. This was quantified as the ratio of percentage CD81 knockout comparing treated versus non-treated cells.

**Figure 2.**
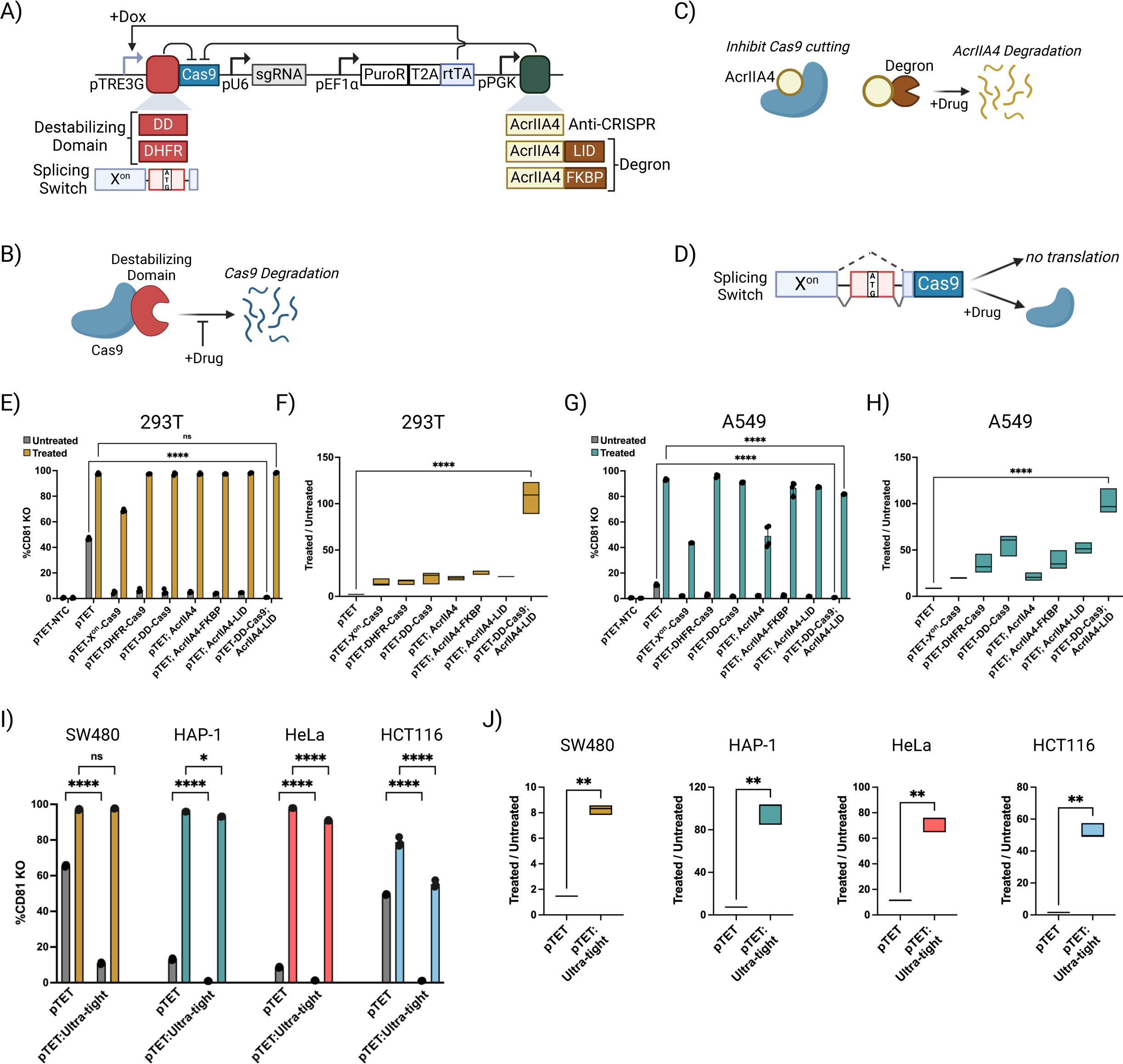
Development of an “ultra-tight” pTET system that is tight and efficient across cell lines. (A) Schematic representation of modifications to the pTET system to minimize leaky Cas9 activity. pTET-NTC line expresses a control guide RNA, while the remaining lines express *CD81* sgRNA. DD: FKBP12^F36VL106P^ destabilizing domain. DHFR: Destabilizing domain based on a mutant *E.coli* dihydrofolate reductase. DD or DHFR fusion degrades leaky Cas9 protein. AcrIIA4: anti-CRISPR gene with an NLS, expressed constitutively from a human PGK promoter. AcrIIA4 inhibits Cas9 activity. The AcrIIA4 gene together with a GSAGSAAGSG flexible linker, is fused to a ligand inducible degradation tag (AcrIIA4-LID) or the FKBP^F36V^ degron (AcrIIA4-FKBP). X^on^ splicing switch: splicing cassette from *SF3B3* with the translation initiation codon in the alternatively spliced exon. (B) Cartoon depicting conditional stabilization of DD or DHFR fused Cas9 in untreated cells. When Cas9 expression is desired, addition of a stabilizing ligand restores protein stability. (C) Cartoon illustrating the strategy to control leaky Cas9 activity by inhibition with AcrIIA4 anti-CRISPR protein. Addition of a ligand specific for each degron tag degrades the anti-CRISPR protein relieving inhibition of Cas9 activity. (D) Schematic representation showing splicing switch mechanism to control leaky Cas9 levels. In untreated cells, exon skipping leads to a Cas9 that is not translated due to lack of the ATG start codon. Upon drug treatment an alternative splicing mechanism retains the start codon leading to translation of Cas9. (E-H) The various pTET modified piggyBac constructs were stably transfected into HEK293T (E, F) and A549 (G, H) cells followed by drug treatment and measurement of CD81 levels as described in Figure 1A and Table 1. (E, G) Percentage of CD81-negative cells was determined in untreated (gray bars) and treated cells (colored bars). (F, H) Box plots display the dynamic range calculated as the ratio of percentage knockout in treated vs. untreated cells for each method. (I) pTET and pTET:Ultra-tight constructs were stably integrated into the indicated cell lines followed by measurement of CD81 levels in untreated (gray bars) and drug-treated (colored bars) cells. (J) Box plots display the ratio of percentage CD81 knockout in treated vs. untreated cells with pTET and pTET:Ultra-tight systems for each cell line. Bar graphs display mean values +/− SD from at least three independent experiments. Statistical significance for differences in percentage of CD81 perturbation was determined by two-way ANOVA with Tukey’s multiple comparisons test (E, G, I), and for the dynamic range plots was determined by either ordinary one-way ANOVA (F, H) or unpaired two-tailed *t*-test with Welch’s correction (J). ns, not significant. *p < 0.05, **p < 0.01, ****p < 0.0001.

**Table 1.** Description of each piggyBac transgenic circuit tested in Fig. 2 including the guide RNA expressed, the control elements to minimize leaky editing, and the drugs required to induce Cas9 protein/function. Unless otherwise indicated, the drugs were used at the following concentrations: 2μg/ml Dox; 1μM Shield1; 2μM TMP; 1μM dTag^v^1; 20-150nM Branaplam depending on the cell line.

| Transgene circuit | sgRNA | Control module 1 | Control module 2 | Inducing drug(s) |
| --- | --- | --- | --- | --- |
| pTET-NTC | NTC | - | - | Dox |
| pTET | CD81 | - | - | Dox |
| pTET-DD-Cas9 | CD81 | DD | - | Dox + Shield1 |
| pTET-DHFR-Cas9 | CD81 | DHFR | - | Dox + TMP |
| pTET; AcrIIA4 | CD81 | - | AcrIIA4 | Dox |
| pTET; AcrIIA4-FKBP | CD81 | - | AcrIIA4-FKBP | Dox + dTag<sup>V1</sup> |
| pTET; AcrIIA4-LID | CD81 | - | AcrIIA4-LID | Dox + Shield1 |
| pTET-DD-Cas9; AcrIIA4-LID | CD81 | DD | AcrIIA4-LID | Dox + Shield1 |
| pTET-X<sup>on</sup>-Cas9 | CD81 | X<sup>on</sup> | - | Dox + Branaplam |

Broadly, the re-engineered systems provided a substantial decrease in background editing and, consequently, a general improvement in dynamic range (Fig. 2E-H). Across tested lines, expression of a non-targeting control sgRNA from the pTET-Cas9 vector (pTET-NTC) demonstrated that CD81 levels were unaffected solely by Dox addition (Fig. 2E, G, S2A). For several, basal *CD81* disruption in cell lines with high (293T, Fig. 2E) and moderate (A549, Fig. 2G) leakiness was reduced to levels similar to that seen with pTET-NTC. K562 cells, which already demonstrated a low background similar to pTET-NTC, and high Dox-induced editing with pTET, did not benefit from additional vector modification (Fig. S2A-B). While there was a general improvement in the dynamic range of editing across the tested conditions, relative to pTET, the peak editing efficiency upon induction was variable; detailed below.

Similar to previous studies, we genetically fused conditional destabilizing “degron” domains ^16,17^ to Cas9 within the pTET vector context, which would lead to rapid turnover of leaky Cas9 protein in the absence of Dox and the small molecule used to stabilize each degron (Fig. 2B, Table 1^6^). Here, DD references a double mutant variant of the FKBP12 protein (FKBP12^F36VL106P^) that is continuously degraded until treated with Shield1^16^. DHFR is an *E.coli* dihydrofolate reductase mutant that destabilizes tagged proteins in the absence of Trimethoprim (TMP)^17^. Although the destabilizing domain-tagged Cas9 variants (pTET-DD-Cas9 and pTET-DHFR-Cas9) displayed comparable percentage knockout in the ON state, background editing was lower with pTET-DD-Cas9 versus pTET-DHFR-Cas9. This resulted in higher dynamic range of basal versus induced activity with the DD-tagged Cas9 module (Fig. 2F, H).

The discovery of anti-CRISPR proteins, several of which can function as natural inhibitors of Cas9 activity, has inspired the development of systems for titratable Cas9 deactivation ^2,18–20^. To investigate the potential application of anti-CRISPRs for reducing basal editing, we modified the pTET vector to constitutively express engineered variants of AcrIIA4, previously identified as a highly effective inhibitor of SpCas9 ^18,21^. For this, we assessed AcrIIA4 fused to a Nucleoplasmin NLS and, as an additional layer of control, tested C-terminal fusions of FKBP12^F36V^ or LID conditional degrons (Fig. 2C). FKBP12^F36V^ (hereafter referred to as FKBP) can be used for selective degradation upon treatment with an appropriate dTag ligand ^22,23^. LID is an engineered variant of FKBP with a cryptic degron at the C-terminus, and is degraded in response to treatment with Shield1^24^. Co-treatment with Dox and dTag (i.e. dTag^V1^) or Shield1 is expected to induce Cas9 expression and in parallel degrade the AcrIIA4, significantly shifting the stoichiometric balance in favor of Cas9 protein/function (Fig. 2C, Table 1).

Accordingly, in the OFF state, continuous expression of non-degradable AcrIIA4 (pTET;AcrIIA4) virtually abolished basal editing (Fig. 2E, G). However, Dox-induced targeting was markedly compromised with this variant in some cellular contexts (see A549 cells, Fig. 2G). Presumably, the induced levels of Cas9 are insufficient to overcome the suppressive effects of the constitutively-expressed anti-CRISPR protein in these contexts. Consistent with this notion, editing efficiency was largely restored and/or consistently high in pTET;AcrIIA4-FKBP and pTET;AcrIIA4-LID, the degron controlled AcrIIA4 lines. In HEK293T and A549 cells, there was no significant difference in drug-induced *CD81* disruption between the two degron-tagged AcrIIA4 systems (Fig. 2E, G). In K562 cells, despite a ∼10% higher editing efficiency with pTET;AcrIIA4-FKBP compared to pTET;AcrIIA4-LID, the dynamic range was similar due to marginally-greater background gene disruption with the AcrIIA4-FKBP fusion (Fig. S2A-B). While subtle differences were observed across cellular contexts for pTET-AcrIIA4-FKBP and pTET-AcrIIA4-LID, the directionality was inconsistent, and we determined these systems to be functionally-interchangeable with one another and the pTET-DD-Cas9 variant.

Targeted degradation and direct inhibition of Cas9 protein both proved to be effective for reducing basal gene editing; we next evaluated whether we could achieve similar results through conditional translation of Cas9. For this, we exploited the “X^on^” alternative splicing switch ^25^ based on a modified, Branaplam-sensitive splicing cassette from *SF3B3*. Placed within the pTET vector (pTET-X^on^-Cas9), immediately downstream of the TRE3G promoter, the modified cassette positions a small exon with an initiator (AUG) codon within an *SF3B3* intron segment. In the absence of Branaplam, the exon with the initiator codon will be spliced over (i.e. excluded) and Cas9 translation will be impaired (Fig. 2D). Upon co-treatment with Dox and Branaplam, a switch in the splicing mechanism will retain the AUG-containing exon, allowing in-frame and full-length production of Cas9 protein. Across all tested cell lines, basal editing was reduced with pTET-X^on^-Cas9 relative to pTET, but this came at a severe cost to perturbation once activated (Fig. 2E, G, S2A). These results suggested that additional measures were needed to optimize the performance and compatibility of the X^on^ and pTET systems.

Encouraged by the targeting efficacy after induction with either the degradable Cas9 or anti-CRISPR control modules, and the potential for further reduction in basal editing, we generated a composite pTET vector with both the DD-Cas9 and AcrIIA4-LID systems (pTET-DD-Cas9;AcrIIA4-LID). Using this design, co-treatment with Dox and Shield1 will induce and stabilize Cas9, while simultaneously relieving inhibition of Cas9 activity through depletion of AcrIIA4. We favored this specific combination of control modules, since it restricts the number of inducing drugs to two rather than the three required for incorporation of either DHFR-Cas9 or AcrIIA4-FKBP. Application of pTET-DD-Cas9;AcrIIA4-LID (hereafter, pTET:Ultra-tight) resulted in a >100-fold dynamic range when comparing the ON versus OFF states for CD81 depletion in both the HEK293T and A549 backgrounds, which was 2-fold greater than any other system in these contexts, with a trend towards improved dynamic range in K562 cells (Fig. 2F, 2H, S2A-B). This was due to the extremely low background editing level, which was indistinguishable from the pTET-NTC control. Induced editing remained high (>80% within polyclonal cell populations) with pTET:Ultra-tight, but with a marginal (∼10%) decrease relative to pTET in both the A549 and K562 cells. We therefore decided to prioritize and expand our evaluation of the pTET:Ultra-tight system due to its unique capacity for nearly eliminating background activity with minimal impact on induced editing.

### Consistent pTET:Ultra-tight functionality across a diverse cell line panel

In order to assess the broad applicability of pTET:Ultra-tight across an expanded set of cell lines, we stably integrated pTET and pTET:Ultra-tight into SW480, HeLa, HAP1, and HCT116. In our initial evaluation of pTET, these four cell lines exhibited a range of basal editing from ∼60% to 8% (Fig. 1B). Stable, polyclonal populations were engineered with pTET or pTET:Ultra-tight and treated with either Dox (pTET) or Dox + Shield1 (pTET:Ultra-tight) for 10 days followed by flow cytometry to measure CD81 levels. Consistent with our previous observations, the pTET:Ultra-tight lines displayed substantially-reduced basal editing in untreated cells relative to pTET in all cell types (Fig. 2I). Of the tested lines, SW480 cells represented an outlying context, where pTET:Ultra-tight was unable to fully reduce background editing. We therefore decided to compare pTET:Ultra-tight to an expanded set of genetic circuits (Fig. S2C), revealing that it remained the superior option for balancing on versus off-state editing (Fig. S2D). Upon drug exposure, the pTET:Ultra-tight system maintained a knockout efficiency greater than 90% in all but one line; HCT116 (∼58%). HCT116 cells also displayed the lowest induced knockout rate (∼80%) tested across all lines with the pTET circuit, suggesting the Dox-regulated circuit requires further optimization within this particular cellular context (Fig. 1B and Fig. 2I). Despite a reduction in knockout efficiency within HCT116 cells with the pTET:Ultra-tight system, the gains in dynamic range between the ON and OFF states, due to a near-complete elimination of background editing, were substantial (>50-fold relative to pTET); a trend that extended to the additional lines tested in this study (Fig. 2J).

### The pTET:Ultra-tight system maintains tight regulation at additional target genes

To investigate whether the pTET:Ultra-tight system maintains stringent regulation at target loci other than *CD81*, we replaced the *CD81* sgRNA in pTET and pTET:Ultra-tight constructs with sgRNAs targeting an expanded cohort of non-essential cell surface markers; β2-microglobulin (B2M), CD298, and CD9. As previously, we generated stable, polyclonal HEK293T cell lines with these constructs, treated each line with either Dox or Dox + Shield1, and evaluated target protein levels 10 days later by antibody staining and flow cytometry. Untreated cells were included as a control to determine basal editing. Consistent with our observations for *CD81*, we detected background editing in untreated cells at all three target loci with the pTET system (Fig. 3A-C). In contrast, pTET:Ultra-tight displayed strikingly-diminished basal editing for each target. Importantly, upon drug exposure, population-wide knockout efficiencies >90% were observed for all target genes. Significant improvement in dynamic range was observed across targets with the pTET:Ultra-tight system (Fig. 3D-F).

**Figure 3.**
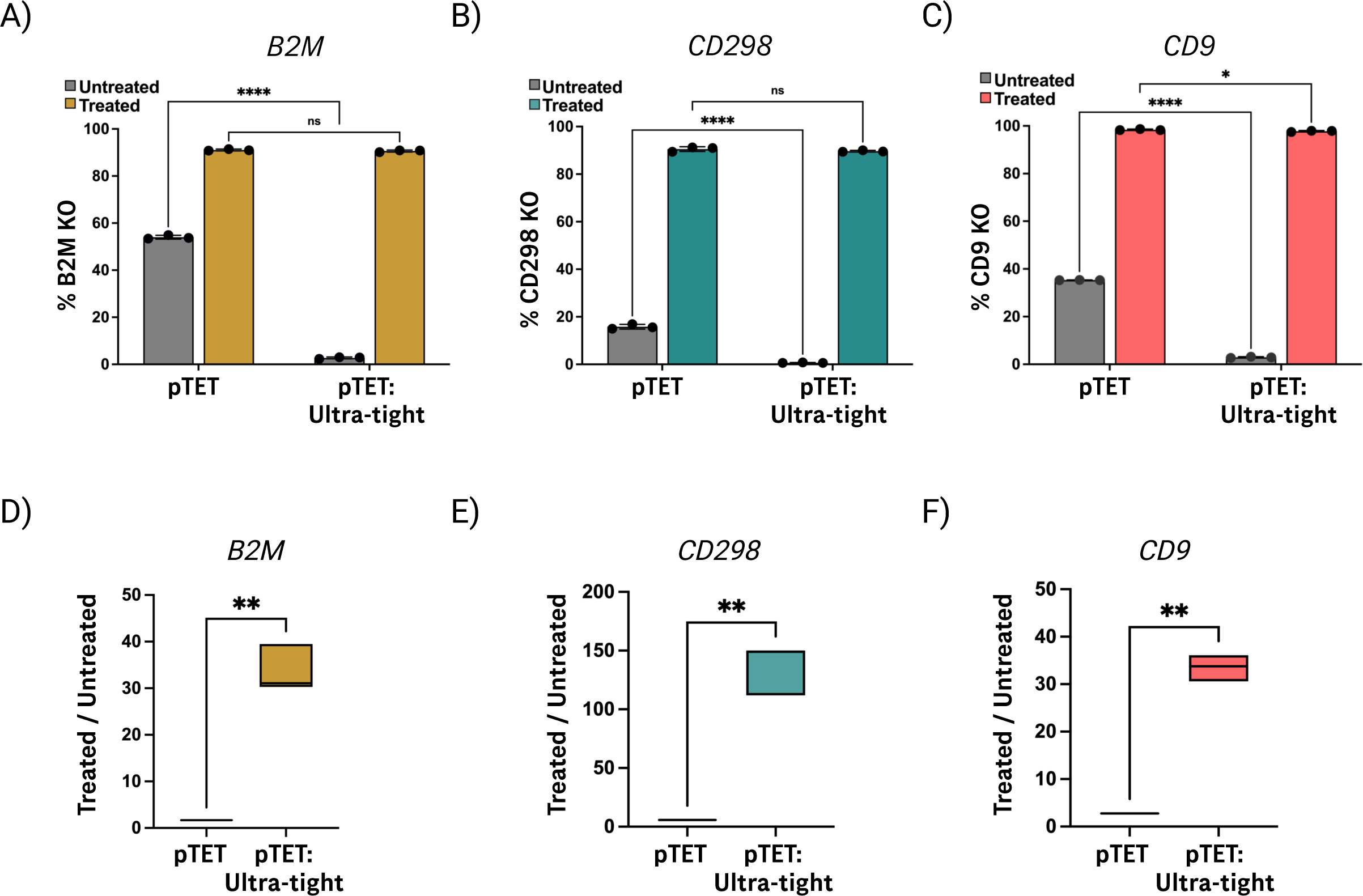
“Ultra-tight” system outperforms pTET at additional target loci. pTET or pTET:Ultra-tight constructs expressing sgRNA targeting B2M (A), CD298 (B) or CD9 (C) were stably integrated into HEK293T cells. Stable cells were left untreated or treated for 10 days with 2μg/ml Dox (pTET) or 2μg/ml Dox + 1μM Shield1 (pTET:Ultra-tight) to induce Cas9 expression. Percentage of CD81 negative cells was measured as described in Figure 1A. (D-F) Box plots display ratio of percentage knockout comparing treated vs. untreated cells for pTET and pTET:Ultra-tight at each target locus. Bar graphs are shown as mean values +/− SD from triplicate experiments. Statistical comparison between the methods was performed with either Two-way ANOVA with Tukey’s multiple comparisons test (A-C) or unpaired two-tailed *t*-test with Welch’s correction (D-F). ns, not significant. *p < 0.05, **p < 0.01, ****p < 0.0001.

### pTET:Ultra-tight system exhibits tight control over essential gene knockout

Large-scale functional genomic screening campaigns have revealed a host of core and condition-specific essential genes ^26^. However, mechanistic study of these genes in a relevant cellular context presents challenges, as loss-of-function leads to a decrease in cell growth or viability and, consequently, elimination of these cells from the overall population. The use of inducible CRISPR/Cas9 would enable study into the temporal progression of phenotypic, transcriptomic, or proteomic changes that take place as cells respond to depletion of these essential factors^27^. However, a leaky system can result in a suboptimal phenotype due to premature disruption of the target and steady, asynchronous decrease in fitness. Although clonal selection can overcome this issue, it is time consuming, laborious, and can misrepresent natural phenotypic or genetic heterogeneity within the population ^28–30^. An ideal inducible system should exercise strict control over target perturbation resulting in a clear-cut phenotype at the population level.

Given the near-absent leakiness and potent induction observed with the pTET:Ultra-tight system, we sought to evaluate its potential for studying an essential gene phenotype. We constructed pTET and pTET:Ultra-tight all-in-one plasmids expressing an sgRNA targeting the Polo-like kinase 1 (*PLK1*) gene. PLK1 is a serine/threonine kinase and a master regulator of cell cycle and mitosis. Disruption of *PLK1* results in proliferation defects. *PLK1* is overexpressed in several cancers and is considered a potential target for cancer therapy ^31,32^. Each of the *PLK1*-targeting constructs, as well as a pTET vector expressing a NTC sgRNA, were stably integrated into HEK293T cells to create polyclonal populations. HEK293T cells were chosen based on the relatively high rate of background editing with pTET (Fig. 1B). Each cell population was either treated with Dox (pTET) or Dox + Shield1 (pTET:Ultra-tight) to induce Cas9 activity or left untreated as a control for background (leaky) targeting. Cell growth was monitored by live imaging over five days in the presence/absence of drugs (Fig 4A). Stable lines were successfully generated for all constructs, but there were notable differences in growth rates in the presence and absence of drug treatment. In the untreated state, the pTET line targeting *PLK1* exhibited reduced proliferation, indicating knockout of the target gene in the absence of induction (Fig. 4B). Conversely, the growth rate of the pTET:Ultra-tight line expressing *PLK1* sgRNA was comparable to the pTET-NTC control line (Fig. 4B). Upon drug treatment, growth was almost fully inhibited in both pTET and pTET:Ultra-tight lines, although the latter demonstrated a greater reduction in growth rate. As previously shown with non-essential gene disruption (Fig. 2E-H, Fig. 3A-F), the pTET:Ultra-tight system demonstrated a significantly-improved dynamic range between treated and untreated states, as compared to pTET (Fig. 4C) While a population-level phenotype was observed with both pTET and pTET:Ultra-tight, we interpret the difference in dynamic range between the systems to reflect “pruning” of cells with leaky and, likely, the greatest capacity for *PLK1* disruption in the pTET context. In that scenario, cells with diminished Cas9 activity would survive but induction would result in a less penetrant phenotype. Our data reinforce the notion that the pTET:Ultra-tight system supersedes pTET for implementing effective, population-level editing control across multiple cell lines and targets including essential genes.

**Figure 4.**
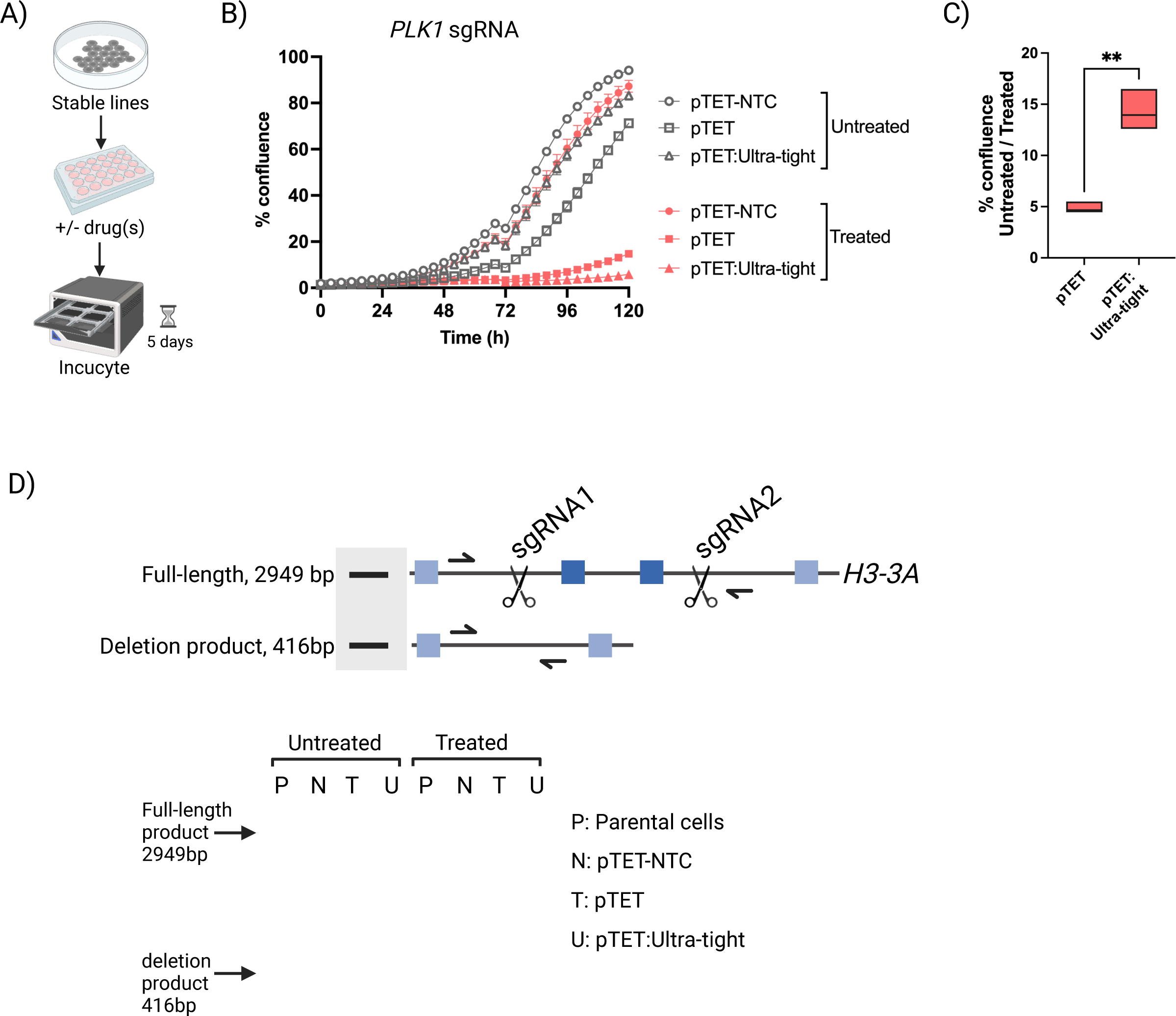
pTET:Ultra-tight system exhibits tight control over essential gene knockout and deletion of a ∼2.9kb genomic region. (A) Schematic overview of the cell growth assay workflow. Stable lines were generated in HEK293T with pTET and pTET:Ultra-tight constructs expressing a *PLK1* guide RNA, while pTET-NTC expresses a control guide RNA. An equal number of cells were plated with or without drug treatment and imaged in an Incucyte over 5 days in 4 h intervals. (B) Growth curves are shown for each line in the presence and absence of drug treatment. Cells were treated with 2μg/ml Dox (pTET and pTET-NTC), 2μg/ml Dox + 1μM Shield1 (pTET:Ultra-tight), or left untreated. Data are shown as mean values ±SEM of triplicate experiments. (C) Box plot displays change in percentage confluence for each line at 120h. Statistical significance was determined with unpaired two-tailed *t*-test with Welch’s correction. **p < 0.01. (D) Top: Schematic of the human *H3-3A* locus showing the sgRNA cut sites and expected amplicon products from wild-type and deleted alleles. Filled blue squares represent exons with exons 2 and 3 depicted as dark blue squares. Half arrows represent PCR primers. Bottom: PCR gel image of amplicon products from the indicated cell lines are shown. Cells were either left untreated or treated with 2μg/ml Dox (parental, pTET, pTET-NTC) or 2μg/ml Dox + 1μM Shield1 (pTET:Ultra-tight) for 72 h.

### Conditional deletion of a ∼2.5kb genomic locus with paired guide RNAs

Thus far, we have explored the general capability of pTET:Ultra-tight for conditional disruption of genes using individual, coding exon-targeting sgRNAs. While effective for creating indels within unique, protein coding segments of the genome, expression of single sgRNAs limits the overall target space. Another option is to co-express a pair of sgRNAs, each targeting opposite ends of a genomic segment, to precisely delete the intervening sequence. This paired sgRNA approach enables excision of complex regulatory DNA elements (i.e. enhancers), non-coding RNAs, repetitive DNA, and genes with family members or pseudogenes sharing highly similar exonic sequences ^33–36^. Furthermore, paired sgRNA expression can be applied for removal of specific exons within a gene or the entire gene ^37^. Genetic loci that cannot be easily perturbed with single sgRNAs may have essential cellular functions that could be revealed or more effectively studied through conditional excision. To these points, we assessed the potential for pTET:Ultra-tight and paired sgRNA expression to induce targeted genomic deletions.

As proof-of-concept for conditional genomic excision with pTET:Ultra-tight, we focused on the *H3-3A* locus, which expresses the non-canonical H3.3A histone variant. *H3-3A* and the near-identical *H3-3B* genes present a distinct use case for targeted deletion, since these share 100% nucleotide similarity across their coding sequences, while the intron and untranslated regions are unique ^38,39^. In addition, *H3-3A* has several pseudogenes with >90% sequence overlap with the *H3-3A* open reading frame ^40^. These uncommon features preclude the use of single sgRNAs for disruption of *H3-3A* via exonic targeting. However, taking advantage of the non-duplicated genomic segments, we designed a pair of sgRNAs targeting intronic sequences flanking exons 2 and 3, which span ∼2.5 kilobases (kb), of the *H3-3A* locus (Fig. 4D).

For expression of the *H3-3A*-targeting sgRNA pair, we cloned one guide within the original U6 promoter cassette in our pTET and pTET:Ultra-tight vectors, while the second guide of the pair was cloned downstream of a newly-integrated H1 promoter. Stable, polyclonal lines were generated in HEK293T with both the pTET and pTET:Ultra-tight constructs, and cells were either left untreated or exposed to inducing drug(s) for 72 hours. After this period we collected genomic DNA and performed PCR with primers spanning the target sites. An amplicon size of ∼2.9 kb indicated presence of the wild-type allele, while successful deletion of the target locus would result in a shorter amplicon of ∼0.4 kb.

In the absence of drug treatment, the ∼2.9 kb product corresponding to the wild-type allele was detected in parental, pTET-NTC control, pTET and pTET:Ultra-tight cells (Fig. 4D). However, only within the pTET cells did we observe the ∼0.4 kb deletion product, indicating leaky editing in this context. Upon drug treatment, the deletion product was observed for both pTET and pTET:Ultra-tight cell lines with a concomitant reduction in signal for the full-length amplicon. Taken together, these results validate the use of pTET:Ultra-tight and paired sgRNA expression for stringent, conditional deletion of genomic loci.

### Development of an enhanced, Dox-independent, conditional CRISPR/Cas9 system

While the Tet/Dox system is commonly employed for conditional transgene expression, it is not uniformly compatible with every cell model or experimental situation. For instance, Dox can disrupt mitochondrial function, making the Tet/Dox system less desirable for studies involving this organelle ^41,42^. Separately, orthogonal control of distinct genetic circuits, to direct or understand complex cellular processes, may be enabled through the use of non-Tet/Dox systems for modulating CRISPR/Cas9 activity^43^. Furthermore, the ability to orthogonally control two processes in the same cell model is a common need that necessitates exploration of an alternative inducible system for controlling CRISPR activity ^44,45^.

Previous studies have applied constitutive Cas9 expression in combination with conditional degradation or alternative splicing for drug-regulated gene editing ^8,46,47^. To evaluate the potency and background editing of Dox-independent induction of Cas9, we designed all-in-one piggyBac vectors with Cas9 expression driven by a constitutive promoter (CMV) and protein levels controlled by either DD or DHFR fusion domains or an X^on^ cassette (Fig S3A). In addition, these vectors contained a constitutive *CD81*-targeting sgRNA expression cassette. Stable, polyclonal HEK293T cell lines were generated for each construct. After antibiotic selection, cell populations were cultured for 10 days with or without respective drug treatments. DD-(N or C-terminal) and DHFR-fused Cas9 expression resulted in 100% depletion of CD81 in the presence or absence of drug treatment, indicating a complete lack of Cas9 activity control under these conditions (Fig S3B). By contrast, the X^on^-Cas9 construct enabled inducible control of Cas9 activity, with ∼88% disruption of *CD81* in the presence of Branaplam. However, this system displayed nearly 20% basal editing in the untreated HEK293T cell context, which encouraged further optimization (Fig S3B).

Based on our ability to mitigate leaky Cas9 activity within the pTET system, we deployed similar strategies to improve upon the X^on^-Cas9 (Fig 5A). To begin, we added the DD or DHFR domain to the N- or C-terminus of Cas9 under control of X^on^. Our data with the optimized TET systems suggested that N-terminal fusion of DD and DHFR would reduce background editing activity (Fig. 2E, G). However, in the context of the X^on^ switch, an N-terminal degron fusion led to 100% leakiness of Cas9 activity (Fig S3C). We reasoned that this could be due to an internal start codon provided by the DD or DHFR domains, which would enable Cas9 translation without Branaplam-induced exon inclusion (Fig S3D). Consistent with this, C-terminal DD or DHFR domain fusions demonstrated a substantial reduction in background effects without impacting ON state editing (Fig 5B). We next tested the addition of an anti-CRISPR protein AcrIIA4 with or without the FKBP degron. The addition of AcrIIA4 dramatically reduced the leakiness of X^on^-Cas9 in HEK293T cells, but at the cost of significantly compromised inducible cutting (Fig 5B). By contrast, inclusion of the AcrIIA4-FKBP control module not only reduced the leakiness but also maintained full ON state editing, offering the best dynamic range of all conditions tested in this cell type (Fig 5B, 5C).

**Figure 5.**
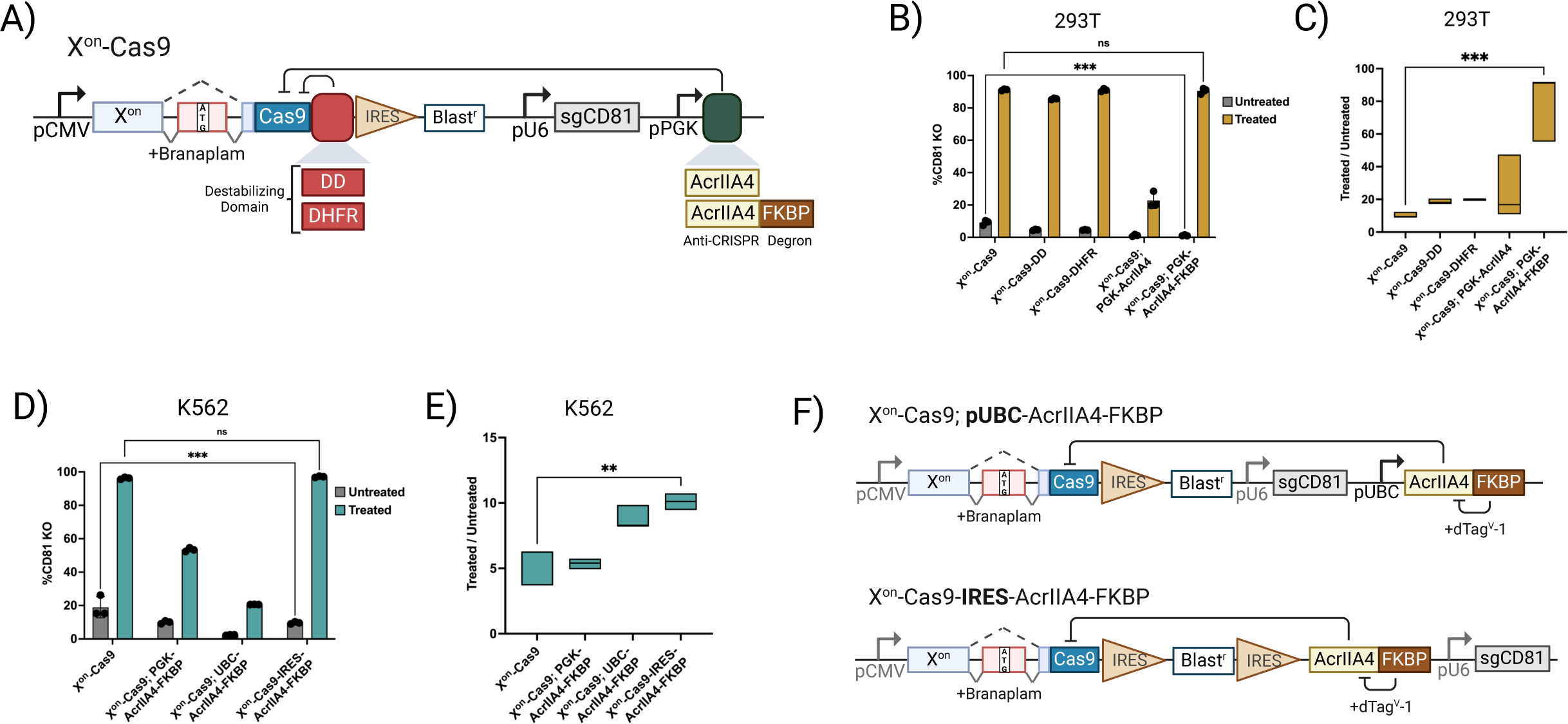
Development of an enhanced Dox-independent inducible Cas9. (A) Schematic diagram of the X^on^ based ultra-tight inducible Cas9. Destabilizing domain DD or DHFR was fused to the C-terminus of Cas9 to reduce the basal expression from the splicing switch. Alternatively, anti-CRISPR proteins with or without the FKBP degron were expressed from a human PGK promoter to inhibit Cas9 activity. (B) *CD81* knockout efficiency was determined in the untreated (gray bars) and treated (colored bars). HEK293T cells with the various versions of X^on^ based inducible Cas9. (C) Box plots show the dynamic range as the ratio of percentage CD81 disruption in treated vs. untreated cells for each method. (D) Bar graphs display *CD81* perturbation efficiency in untreated (gray bars) and treated (colored bars) K562 cells stably integrated with the different versions of X^on^ based inducible Cas9. (E) Box plots display the dynamic range for each method. (F) Schematic diagram of the two additional versions to control the anti-CRISPR module in the X^on^ system. Bar graphs are presented as mean values +/− SD from triplicate experiments. Statistical significance was determined with either Two-way ANOVA with Tukey’s multiple comparisons test (B,D) or ordinary one-way ANOVA (C, E). ns, not significant. **p < 0.01, ***p < 0.001.

We then tested the X^on^-Cas9;AcrIIA4-FKBP combination in K562 cells. Although background editing was nearly absent in this cell line with the pTET system, the unmodified X^on^-Cas9 system displayed ∼19% leaky target disruption (Fig 5D). Use of the X^on^-Cas9;AcrIIA4-FKBP vector reduced leakiness in this cellular context, but it also resulted in poor induction of Cas9 activity (Fig 5D, E). Within this vector design, AcrIIA4 expression is under the control of a PGK promoter (Fig 5A). Separate studies have observed slightly higher or more stable transgene expression from the PGK promoter relative to other constitutive promoters, albeit with context-dependent results ^48,49^. Therefore, we hypothesized that the PGK-driven levels of AcrIIA4 protein were too high, even with dTag-mediated degradation, to fully reverse inhibition of Cas9 function, and that weaker AcrIIA4 expression was needed. To this, we evaluated two distinct strategies aimed at lowering the baseline expression of AcrIIA4 such that it would maintain suppression of Cas9 in the uninduced state but allow for more complete reversal of this effect when cultured with a dTag molecule. This included exchange of the PGK promoter for the generally less-expressive UBC promoter ^48,49^ or construction of an extended, polycistronic transgene with AcrIIA4 expressed by way of an IRES downstream of the CMV-X^on^-Cas9 cassette (Fig 5F). IRES elements are commonly used to express multiple genes simultaneously and have been reported to drive lower expression of the downstream gene compared to the upstream, cap-dependent translated segment ^50^. Both background and induced editing were greatly reduced with the UBC promoter driving the anti-CRISPR module suggesting that UBC is stronger than PGK in our tested K562 cell line (Fig 5D). By contrast, IRES-AcrIIA4-FKBP led to significantly-reduced leakiness with no compromise in induced cutting and displayed the best dynamic range of editing (Fig 5D, 5E). Together, these results demonstrate that non-Dox-inducible Cas9 expression systems can enable high efficiency and low background editing, aided through co-expression of degradable anti-CRISPR proteins, but cell type-dependent variability necessitates empirical testing for optimal performance.

### An enhanced, inducible CRISPRi system for potent, tightly regulated gene silencing

The use of Cas9-mediated dsDNA breaks for gene perturbation may not be suitable for certain applications^51^. Accordingly, CRISPR interference (CRISPRi) provides a complementary and powerful tool for loss-of-function purposes. Here, a nuclease-dead Cas9 (dCas9) is fused to a Krüppel-associated box (KRAB) repressor or various chromatin and DNA modifying domains to facilitate transcription silencing of an intended target gene through a promoter or enhancer-targeting sgRNA ^52,53^. While dCas9 fused to the *KOX1* KRAB domain has commonly been used for CRISPRi, *ZIM3* KRAB has emerged as a more potent alternative ^54,55^, in both constitutive and inducible vector contexts ^54,56,57^. Recognizing the utility of inducible CRISPRi for targeting both coding and non-coding regions, we sought to identify key parameters that enable robust, population-wide gene silencing and minimal background through expression of dCas9-KRAB variants.

We began by evaluating a foundational set of TET-inducible CRISPRi systems. For this, KRAB domains from either *KOX1* (pTET-KOX1) or *ZIM3* (pTET-ZIM3) were fused to dCas9 and this cassette was positioned downstream of a TRE3G promoter within a lentiviral backbone with a constitutive blasticidin resistance and rtTA expressing transgene (Fig 6A). Lentiviral infection with this vector was used to generate stable, polyclonal DLD1 (colorectal cancer) cell lines. These lines were subsequently transduced with lentivirus expressing an sgRNA targeting the *CD81* promoter. Each cell population was treated with Dox or left untreated for 7-days and then assessed for CD81 depletion. The pTET-KOX1 cell line exhibited minimal effect on CD81 levels in the absence of Dox, but also correspondingly low (<50%) activity upon induction (Fig 6B, S4A). Conversely, pTET-ZIM3 demonstrated substantial improvement for silencing *CD81* upon Dox induction, although this potency translated to noticeable leakiness, impacting ∼40% of the untreated population (Fig 6B, S4A). This prompted us to optimize the ZIM3-based CRISPRi system to reduce its leakiness and maintain its activity via the principles applied for inducible expression of Cas9 nuclease.

**Figure 6.**
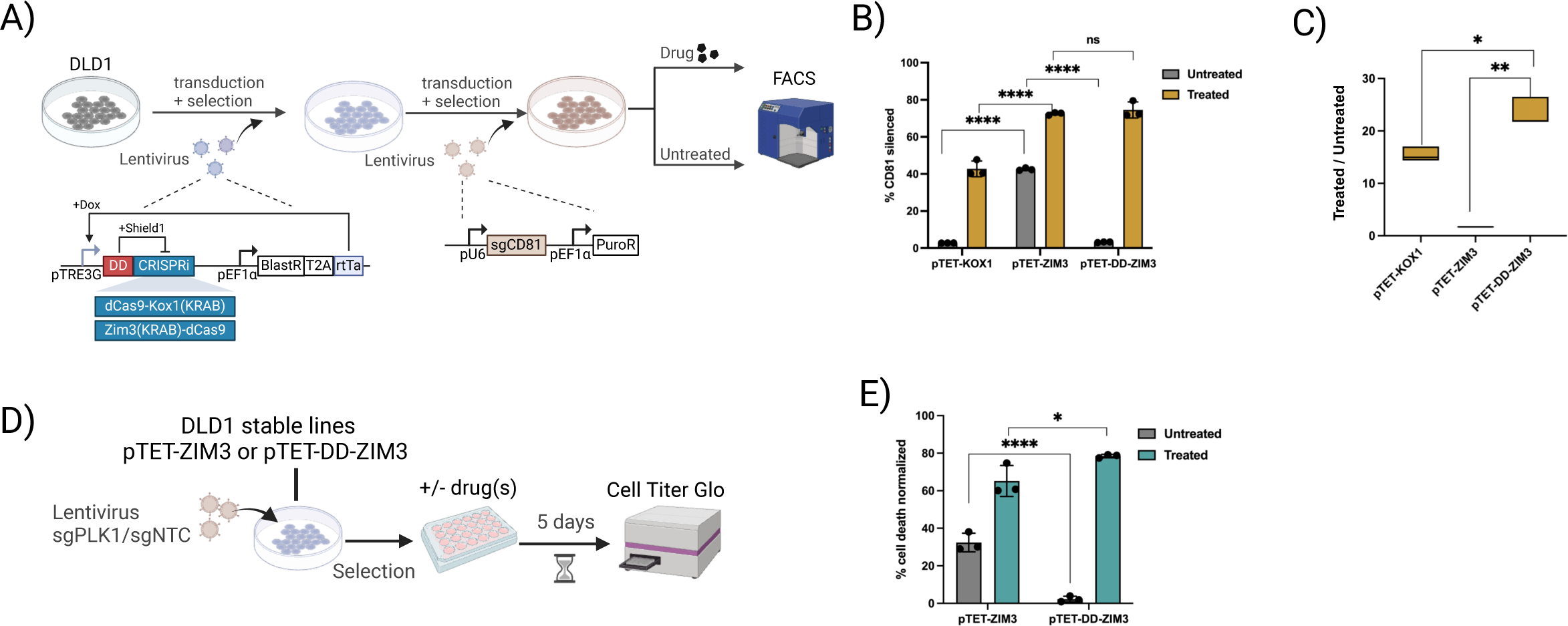
Ultra-tight inducible CRISPRi system. (A) Workflow for examining the inducible CRISPRi system in DLD1 cells. The inducible dCas9KRAB constructs were integrated into the cells via lentivirus transduction. The lentiviral cell line was established first and then the sgRNA was introduced in a second round of transduction. (B) *CD81* silencing in the inducible CRISPRi cells was measured by flow cytometry. pTET-KOX1, Tet-On dCas9 fused with KRAB domain from KOX1; pTET-ZIM3, Tet-On dCas9 fused with KRAB domain from ZIM3; pTET-DD-ZIM3, Tet-On system expressing a fusion protein containing DD, KRAB from ZIM3, and dCas9. CD81 expression was measured after 7 days of treatment or non-treatment. Bar graph displays the percentage of cells with *CD81* silenced. (C) Box plots display CRISPRi dynamic range shown as the ratio of CD81 silencing in induced vs uninduced cells. (D) Workflow for examining *PLK1* silencing in the inducible CRISPRi DLD1 cells. pTET-ZIM3 and pTET-DD-ZIM3 lines from (A) were transduced with lentivirus expressing sgRNA targeting *PLK1* or a non-targeting control. The cells were selected and then treated or untreated for 5 days before measurement by Cell Titer Glo assay. (E) *PLK1* silencing shown as normalized cell death. Normalized cell death is calculated as luminescence measurements of (cells with sgNTC - cells with sg*PLK1*)/(cells with sgNTC) in each condition. Bar graphs are shown as mean values +/− SD from triplicate experiments. Statistical comparison between the methods was performed with Two-way ANOVA with Tukey’s multiple comparisons test (B), uncorrected Fisher’s LSD (E), or RM One-way ANOVA (C). ns, not significant. *p < 0.05, **p < 0.01, ***p < 0.001, ****p < 0.0001.

To improve dynamic range between the CRISPRi ON and OFF states, we incorporated an N-terminal DD tag to the ZIM3-dCas9 cassette, resulting in pTET-DD-ZIM3. In this instance, a separate anti-CRISPR module was not included to avoid complications with viral particle production related to cargo size limitations with lentiviral backbones. Induction of pTET-DD-ZIM3 cells by combined Dox and dTag treatment reduced CD81 levels similar to pTET-ZIM3 with minimal background gene silencing. (Fig. 6B). As a result, dynamic range for the inducible, degron-enabled CRISPRi system was increased ∼13 fold over pTET-ZIM3 (Fig 6C). Similar to our evaluation of pTET:Ultra-tight, we compared pTET-ZIM3 with pTET-DD-ZIM3 for disruption of the essential gene, *PLK1* (Fig 6D). Confirming the improved function of pTET-DD-ZIM3, background editing was significantly reduced, while the strength of phenotype upon induction was elevated related to pTET-ZIM3 (Fig 6E). For CRISPRi, the combination of Dox induction with degron-mediated stabilization of ZIM3-dCas9 offers the most potent tool for gene silencing with minimal background.

## Discussion

The ability to control the timing and dosage of gene expression can be critical for understanding dynamic or cell state-dependent biological processes. Accordingly, drug-inducible CRISPR/Cas9 systems enable on-demand disruption of gene activity within specific conditions that would otherwise be masked or difficult to study when using static targeting methods. Current approaches for inducible Cas9 have revealed unique biological insights, but broadened application of these tools has been limited by a combination of background editing and/or modest gene targeting efficacy in the presence of the inducing agent(s), compared to constitutive disruption systems ^47^. Unintended editing is distinctly problematic for inducible Cas9 nuclease expression. For example, slow and progressive accumulation of DNA damage in the off-state may degrade the cell population and hamper the study of essential genes ^3,4,14^.

To maximize the potency and limit background effects of chemically-inducible gene perturbation, we implemented complementary strategies for regulating Cas9 translation, stability, and function. Through a newly-developed, degradable anti-CRISPR module and multi-layered Cas9 control strategies, we established the pTET:Ultra-tight system. To enable pTET:Ultra-tight for general use, we incorporated these modules into an all-in-one, stably-integrating piggyBac vector. This vector allows for single-step creation of cell lines with the capacity for population-wide, inducible gene targeting and negligible baseline Cas9 nuclease activity without the need for single cell cloning. Comparison of inducible Cas9 formats across cell lines and targets showed consistently-high efficacy, low background, and superior overall performance with pTET:Ultra-tight. Furthermore, we demonstrated the ability of pTET:Ultra-tight to enhance the study of an essential gene (*PLK1*) by eliminating background editing, which can lead to pruning (death or reduced growth) of cells with leaky Cas9 activity. In addition, we demonstrated a unique application for the pTET:Ultra-tight system via co-expression of target-flanked sgRNAs to conditionally-delete *H3-3A*. This gene shares near-identical coding exon sequence with other *H3-3* family members, preventing the use of individual sgRNAs to introduce frame-altering mutations. While we focused on deletion of a protein-coding gene in this instance, this approach can likely be applied across a broad range of genomic elements, including unique exons or untranslated regions (UTRs), non-coding RNAs, and regulatory DNA. Multiplexed expression of sgRNAs (each directed to a distinct gene) from the pTET:Ultra-tight context should also facilitate targeting of multiple genes in the same cell.

Evaluation of pTET:Ultra-tight across cell lines revealed areas for future system optimization. HCT116 cells demonstrated the lowest degree of target disruption (60-80%) in the presence of Dox, with or without the degron and anti-CRISPR control modules. This may be addressable through Dox-sensitized rtTA elements ^58^. Conversely, while pTET:Ultra-tight showed the greatest reduction of background editing of all tested systems in SW480 cells, ∼10% of cells showed uninduced target disruption. To reduce basal Cas9 activity, alternative degron modules or boosted expression of the anti-CRISPR module could prove beneficial. For either circumstance, different modules ^59^ for induced Cas9 transcription may be more dynamic than the TET/Dox system, and these could be considered on a cell type-by-cell type basis for specialized applications.

Although pTET:Ultra-tight proved effective under a range of conditions, its reliance on Dox for induction makes it unsuitable for applications where this drug might impact cell physiology ^60^. Separately, building complex synthetic gene circuits may require the use of Dox-induction for elements other than Cas9, necessitating alternative means of control for gene editing activities. To accommodate these possibilities, we took advantage of the recently developed X^on^ system, which enabled us to develop a module where Cas9 translation could be controlled through a Branaplam-induced alternative splicing mechanism. Despite strong Cas9 induction, resulting in population-wide editing in separate cell line contexts, removal of background editing required additional control measures. Comparison of degron-linked Cas9 to the degradable anti-CRISPR modules, where Cas9 would be induced by co-treatment with Branaplam and either Shield1 or a dTag molecule, showed lower uninduced editing with the anti-CRISPR module. Dosage of the anti-CRISPR was critical, and distinct expression strategies were required across cell lines to increase the ratio of Cas9 activity in the ON versus OFF states. Recently, the X^on^ system has been modified and used to reduce background editing of an inducible Cas9 cassette, albeit without additional modules^61^. We anticipate further evolution of this technology in the coming years, which will only increase the robustness and flexibility of inducible genome manipulation. However, Branaplam and other molecules in this class are expected to trigger alternative splicing of endogenously expressed transcripts ^62^ beyond the SMN intron used for the X^on^ module. This will need to be taken into consideration for select cell contexts and applications.

Lastly, for experiments that benefit from targeted transcriptional repression and the avoidance of dsDNA breaks, we have established a lentiviral system for inducible CRISPRi. For this, we integrated the ZIM3 KRAB domain within a dual Dox + degron-linked dCas9 expression context, resulting in pTET-DD-ZIM3. Combining these modules demonstrated superiority for both induced and background silencing levels compared to Dox + ZIM3-dCas9 alone or a Dox + dCas9-KOX1 KRAB fusion. As with pTET:Ultra-tight, the pTET-DD-ZIM3 provided high dynamic range in a cell viability assay when targeting *PLK1*. Notably, gene silencing by the ZIM3 KRAB domain is expected to be transient. While spurious, low level expression by Cas9 nuclease may result in measurable, progressive gene perturbation, this is less likely to be the case for ZIM3-dCas9^63^. As such, adding a third control module within pTET-DD-ZIM3 (i.e., degradable anti-CRISPR) may offer little improvement in dynamic range and could potentially reduce the quality of virus production or expression of transgenes when using lentivirus, given the limited cargo capacity of the lentiviral genome.

Together, we provide an enhanced gene editing toolbox through the creation of “Ultra-tight” inducible CRISPRn, splice switch control over Cas9 activity, and related CRISPRi systems. These permit population-level gene disruption, often with marginal or no measurable background activity. The consistent and exceptionally-high dynamic range of efficacy for pTET:Ultra-tight was demonstrated in comparison to alternative inducible Cas9 formats across all tested cell types. We simplified and broadened the applicability of pTET:Ultra-tight by integrating it within an all-in-one transgenic vector for single-step cell line production and co-expression of one or more sgRNAs. Importantly, the core components and conceptual framework of our inducible gene perturbation systems, which rely on multi-layered control modules, could likely be adapted to create vectors for other functions, such as base editing, prime editing, or CRISPR activation, or to work with alternative effector proteins including Cas12a, Cas13, and their associated anti-CRISPRs ^19,64^. With continuous innovation in targeted protein degradation ^65,66^ and inducible genetic switch methodology, there will likely be more opportunities to expand the versatility and application of conditional genome engineering technologies ^67,68^.

## Supplementary Data

**Figure S1.**
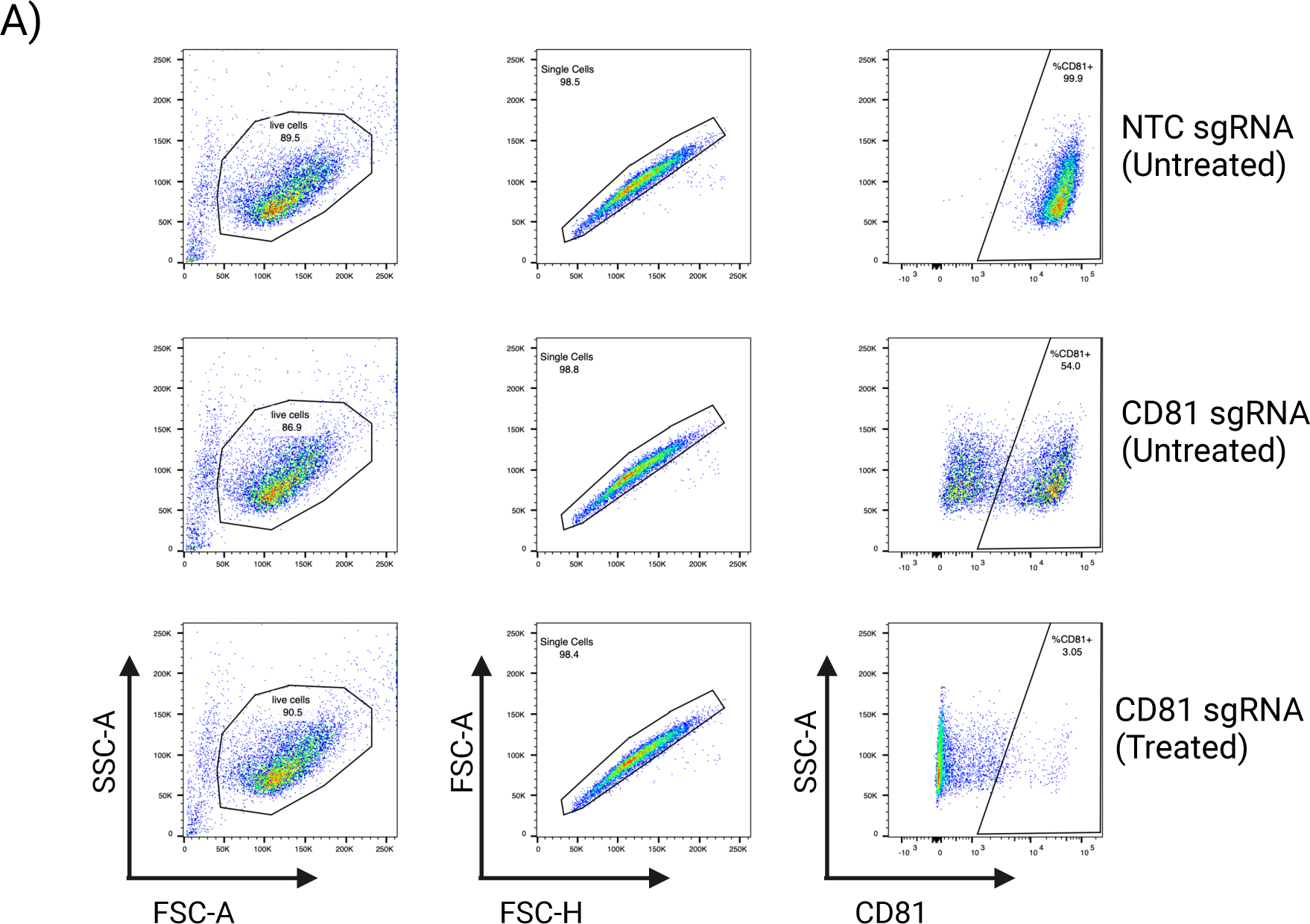
Gating strategy for flow cytometric analysis. (A). Representative flow cytometry plots of pTET and pTET-NTC stable HEK293 cells with and without Dox treatment. SSC-A (side scatter) and FSC-A (forward scatter) profiles were used to isolate the live cell population. Doublets were excluded using FSC-A and FSC-H profiles. The pTET-NTC control line was stained in parallel to define the CD81 positive population.

**Figure S2.**
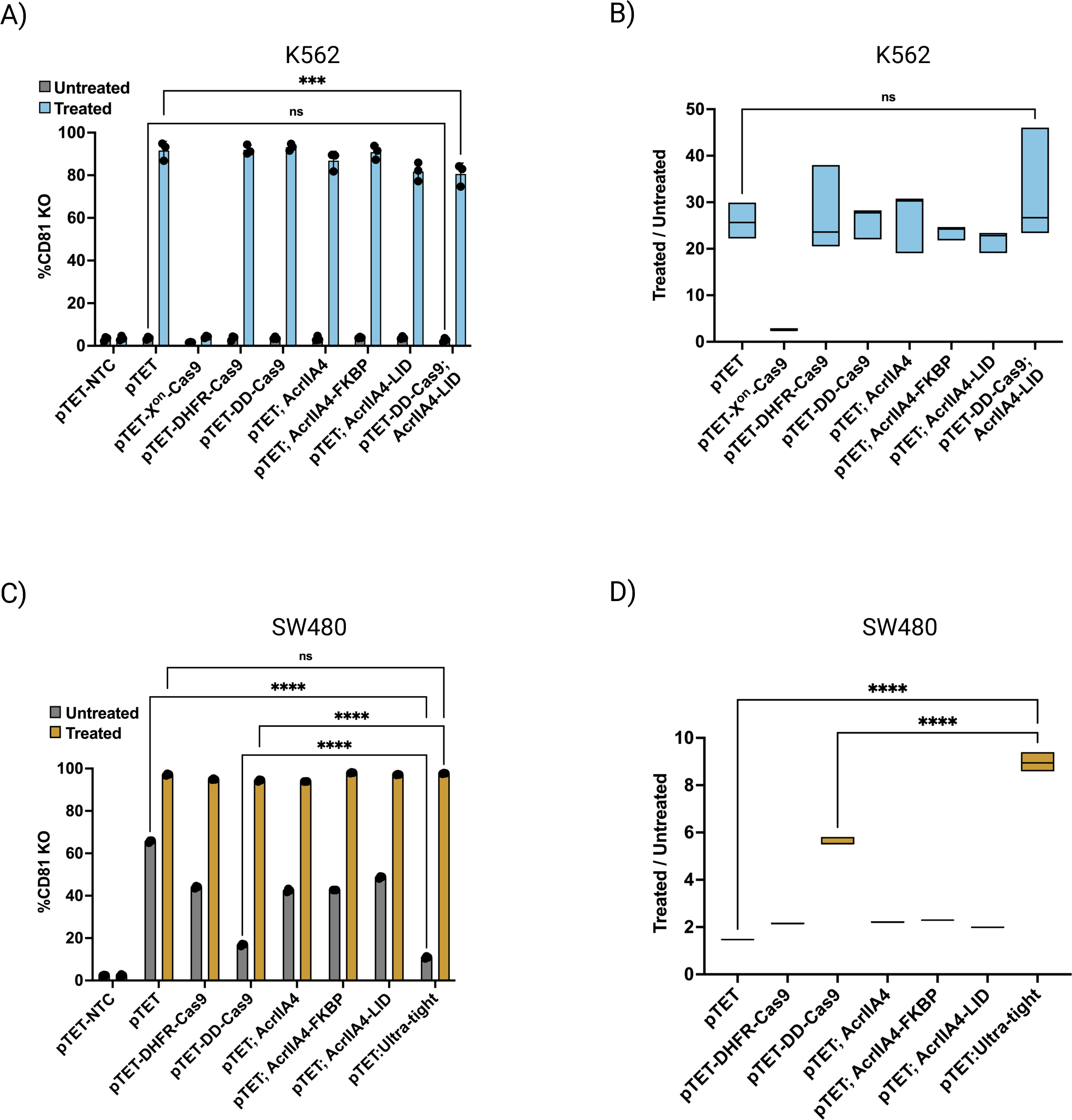
Comparison of the various methods in additional cell lines. The different pTET constructs were stably transfected into (A-B) K562 cells or (C-D) SW480 cells, followed by drug treatment for 10 days and flow cytometry. The percentage of CD81 negative cells in treated (colored bars) and untreated (gray bars) cells is shown. Data are presented as mean values +/− SD from triplicate experiments. (B, D) Box plots display the dynamic range of Cas9 activity calculated as the ratio of percentage knockout in treated vs. untreated cells for each method. Statistical significance was determined by either Two-way ANOVA (A, C) or ordinary one-way ANOVA (B, D) adjusted with Tukey’s multiple comparisons test. ns, not significant. ***p < 0.001, ****p < 0.0001.

**Figure S3.**
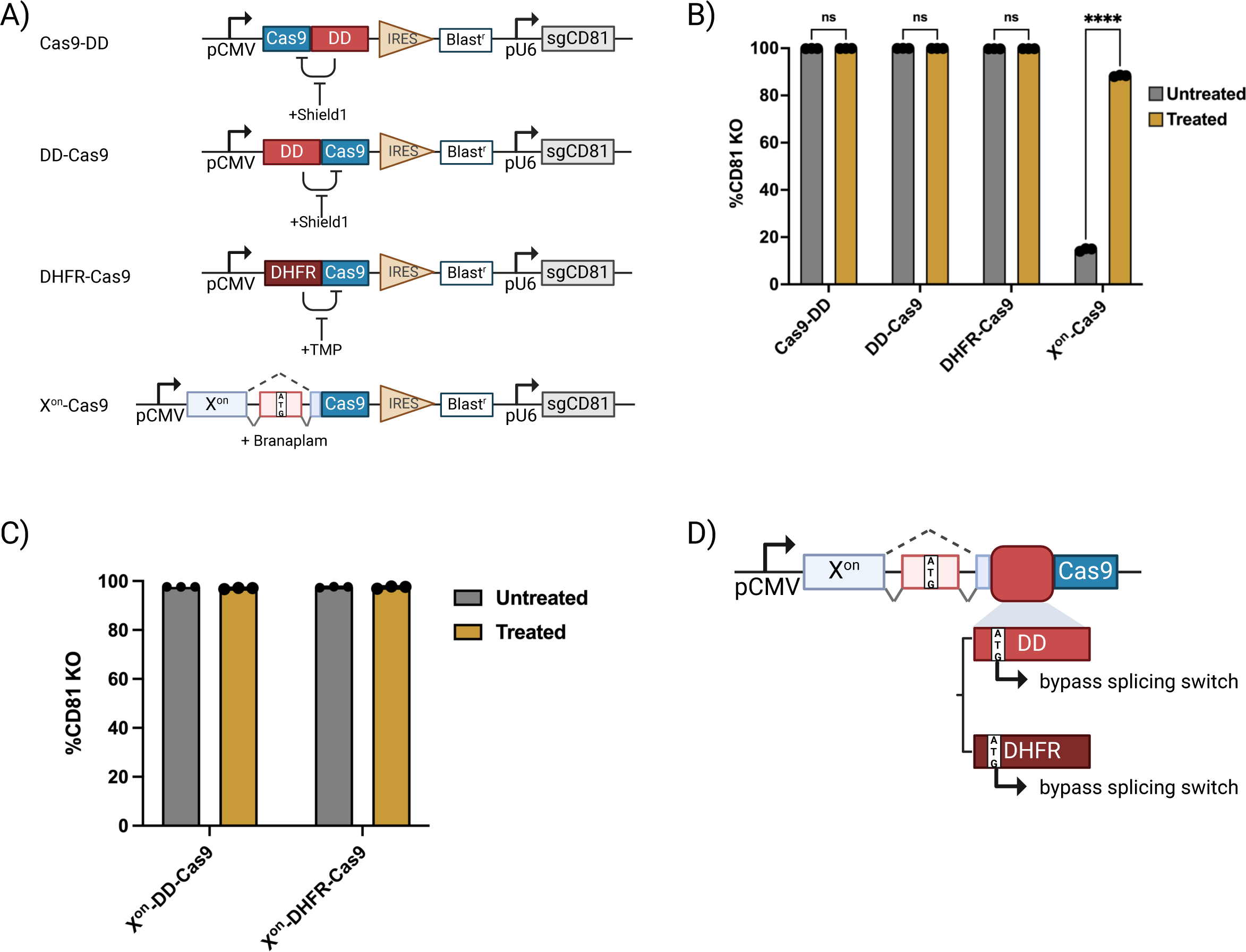
Dox independent inducible Cas9 comparison. (A) Schematic diagram of dox independent inducible Cas9 systems. DD domain or DHFR domain is fused to Cas9 at the N- or C-terminus. In the X^on^ Cas9, ATG is deleted from the Cas9 coding sequence and provided in the alternatively spliced exon. (B) *CD81* knockout percentage in the untreated (gray bars) and treated (colored bars) HEK293T cells that stably express inducible Cas9 with a single regulatory module. (C) *CD81* knockout percentage in the untreated (gray bars) and treated (colored bars) HEK293T cells with DD or DHFR in combination with X^on^ switch. (D) Schematic diagram of the potential mechanism of Cas9 leaky expression observed in (C). Internal ‘ATG’ in the DD or DHFR domain bypassed the requirement of exon 2, rendering the X^on^ Cas9 constantly active. Bar graphs are presented as mean values +/− SD from triplicate experiments. Statistical significance was determined by two-way ANOVA adjusted with Tukey’s multiple comparisons test. ns, not significant. ****p < 0.0001.

**Figure S4.**
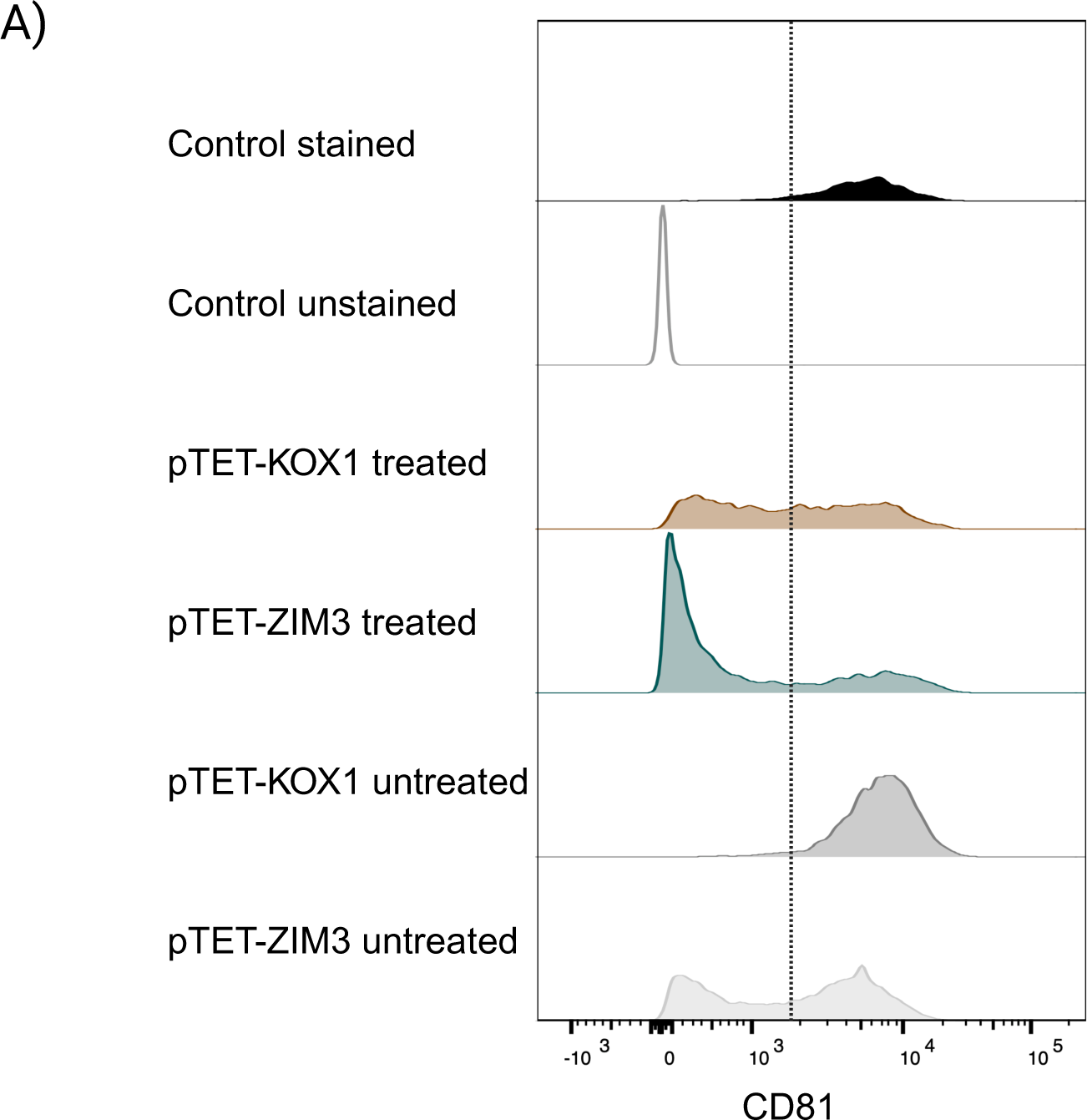
*CD81* silencing with KOX1 vs. ZIM3 based inducible CRISPRi systems. (A) Representative flow cytometry result of the CD81 stained cells with inducible CRISPRi systems. DLD1 cells expressing sg*CD81* and pTET-KOX1 or pTET-ZIM3 were treated with 250 ng/ml Dox or untreated for 7 days. The cells were stained with CD81 antibody conjugated with PE. CD81 positive and negative cells were separated by the dashed line. Control: parental DLD1 cells.

## Materials and Methods

### Cell culture and stable line generation

All parental cell lines were obtained from Genentech cell bank (gCell), validated by STR profiling and tested for mycoplasma. Genome integration was accomplished using the PiggyBac transposition system where each piggybac transposon plasmid was delivered into cells together with a plasmid expressing the piggybac transposase. Cell culture conditions and transfection methods are described in Supplementary information (Table 1). Briefly, cells were seeded as indicated and transfected in 1mL Opti-MEM. Transfection reagents were used as per manufacturer’s recommendations and Opti-MEM was replaced with growth media 4 h post-transfection. Plasmid constructs were generated by gene synthesis (Genscript) and included either Puromycin or Blasticidin resistance markers to select for stable cells. Unless otherwise indicated, stable lines were enriched with 1μg/ml Puromycin or 20μg/ml Blasticidin. Sequences of plasmids used in the study are listed in Supplementary information -Table 5. Guide RNA sequences for CRISPRn and cell culture drugs used in this study are listed in Supplementary information - Tables 2 and 4.

### Flow Cytometry

In brief, adherent cells were lifted by TrypLE at 37 degrees for 5 minutes and resuspended in growth medium. Dissociated adherent cells or suspension cells were spun down at 500g for 5 minutes and resuspended in 100μl FACS buffer (PBS, 0.5% BSA, 0.05% NaN3 sodium azide) with 5μl of antibody. After 10 minute incubation on ice, the cells were washed three times with FACS buffer and passed through a 35-μm cell strainer. Data was collected on BD Symphony instruments using FACSDIVA acquisition software. Data was analyzed by FlowJo v10.10.0. The gating strategy to isolate live and single cell populations is shown in Supplementary Figure 1A. The following antibodies were used for flow cytometric analysis. CD81 Monoclonal Antibody (1D6-CD81) APC (ThermoFisher Scientific, Cat. No. 17-0819-42), CD81 antibody PE (BioLegend, Cat. No. 349505), FITC anti-human β2-microglobulin Antibody (BioLegend, Cat. No. 316304), PE/Cyanine7 anti-human CD9 Antibody (BioLegend, Cat. No. 312115) and PE/Cyanine7 anti-human CD298 Antibody (BioLegend, Cat. No. 341707).

### Lentivirus production and transduction

HEK293T cells were seeded and grown to 80-90% confluency at the time of transfection. Lentiviral plasmids expressing target genes were co-transfected with packaging plasmids pDelta8.9 and pVSVG into HEK293T cells using Lipofectamine 2000 reagent. 6 hours after transfection, the medium containing transfection mix was aspirated and fresh growth media was added to the cells. Lentiviruses were collected twice from the supernatant 24 hours and 48 hours post transfection. Cell debris was removed by passing the crude supernatant through the 45 μm filter. Lentivirus was concentrated by Lenti-X (Takara Bio). DLD1 cells were cultured in RPMI 1640 supplemented with 10% TET-free FBS, 2mM L-Glutamine and pen-strep. Cells were seeded in 6-well Corning plates and grown to 50% confluency on the day of transduction. The lentivirus and 8μg/ml polybrene were added to the DLD1 cells. Cells were spin infected at 1800 rpm for 45 minutes and incubated for 48 hours before passage. DLD1 was first transduced with lentivirus expressing inducible CRISPRi systems and selected with 10 μg/ml Blasticidin to obtain the stable cell line, and then transduced with lentivirus expressing sgRNA and selected with 2 μg/ml puromycin.

### PCR genotyping

Genomic DNA was isolated from untreated and treated cells 72 hours post-treatment using QuickExtract™ DNA Extraction Solution (Lucigen). Both full-length and deleted forms of the target locus were amplified with Q5® High-Fidelity 2X Master Mix (NEB) using primers (forward 5′-GGCTTTGTAGAATGCTCGCTG-3′; reverse 5′-TTTAACGACCCCCAACTGCAT-3′) flanking exons 2 and 3 within the *H3-3A* gene. Amplicons were resolved on an agarose gel and images were obtained using the ChemiDoc MP imaging system (Bio-Rad).

### Cell proliferation assay

An equal number of HEK293T pTET-NTC, pTET or pTET:Ultra-tight stable cells expressing *PLK1* sgRNA, were seeded at a low density in growth media with or without inducing drug(s). Cell growth was monitored using the IncuCyte S3 Live Cell Imaging and Analysis Instrument (Sartorius). Phase contrast cell images were collected using a 10× objective lens within the instrument every 4 h over 5 days and cell proliferation was determined based on confluence.

### Cell Viability Assay

DLD1 stable cells generated by transduction of sgNTC lentivirus or sg*PLK1* lentivirus cocktail *(PLK1* sgRNA1 +*PLK1* sgRNA2) were replated into 96 well format at 1000 cells/well in 80μl medium. The following day, 20 μl of medium with or without treatment (final concentration: 250 ng/mL dox, 250 ng/mL dox + 1μM shield1) was added to begin induction. On day 5 after induction, CellTiter-Glo Luminescent Cell Viability Assay (Promega #G7572) was used to read out cell viability on the EnVision plate reader (PerkinElmer/Revvity). Guide RNA sequences are provided in Supplementary Information - Table 3.

## Supplementary information

**Table 1.** Cell culture and transfection conditions.

| Parental cell line | Growth media | Cell type | PiggyBac transfection |
| --- | --- | --- | --- |
| HEK293T | DMEM + 10% TET-free FBS + 1% NEAA + 2mM L-Glutamine; 1X pen-strep | adherent | Seed $3 \times 10^5$ cells/well (6-well plate) and the next day transfect with Lipofectamine 2000 (ThermoFisher Scientific); 2μg total DNA; 3:1 ratio of pB plasmid:pB transposase. |
| A549 | RPMI 1640 + 10% TET-free FBS + 2mM L-Glutamine; 1X pen-strep | adherent | Seed $5 \times 10^5$ cells/well (6-well plate) and the next day transfect with GeneJuice (Novogene); 2μg total DNA; 10:1 ratio of pB plasmid:pB transposase. |
| SW480 | RPMI 1640 + 10% TET-free FBS + 2mM L-Glutamine; 1X pen-strep | adherent | Seed $5 \times 10^5$ cells/well (6-well plate) and the next day transfect with GeneJuice (Novogene); 2μg total DNA; 3:1 ratio of pB plasmid:pB transposase. |
| HeLa | EMEM + 10% TET-free FBS + 2mM L-Glutamine; 1X pen-strep | adherent | Seed $5 \times 10^5$ cells/well (6-well plate) and the next day transfect with Lipofectamine 3000 (ThermoFisher Scientific); 2μg total DNA; 3:1 ratio of pB plasmid:pB transposase. |
| HAP-1 | IMDM + 10% Tet-free FBS + 2mM L-Glutamine; 1X pen-strep | adherent | Seed $2 \times 10^5$ cells/well (6-well plate) and the next day transfect with GeneJuice (Novogene); 2μg total DNA; 3:1 ratio of pB plasmid:pB transposase. |
| HCT116 | RPMI 1640 + 10% TET-free FBS + 2mM L-Glutamine; 1X pen-strep | adherent | Seed $5 \times 10^5$ cells/well (6-well plate) and the next day transfect with GeneJuice (Novogene); 2μg total DNA; 10:1 ratio of pB plasmid:pB transposase. |
| K562 | RPMI 1640 + 10% TET-free FBS + 2mM L-Glutamine; 1X pen-strep | suspension | Transfect $1 \times 10^6$ cells with GeneJuice (Novogene); 2μg total DNA: 3:1 ratio of pB plasmid : pB transposase. |

**Table 2.**
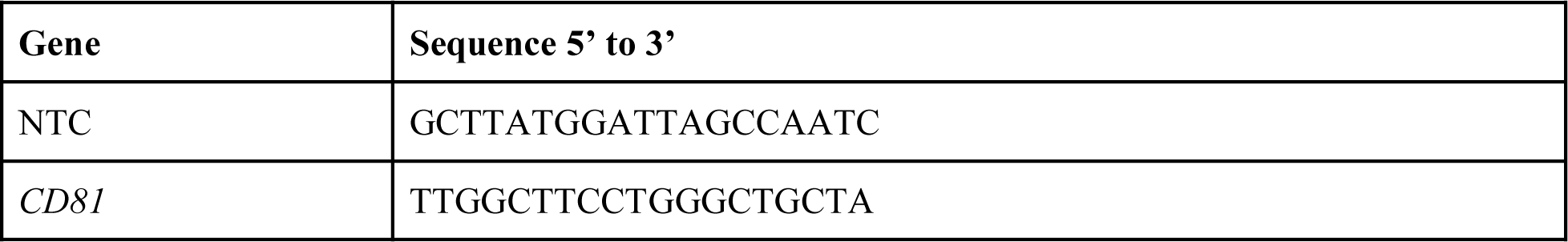
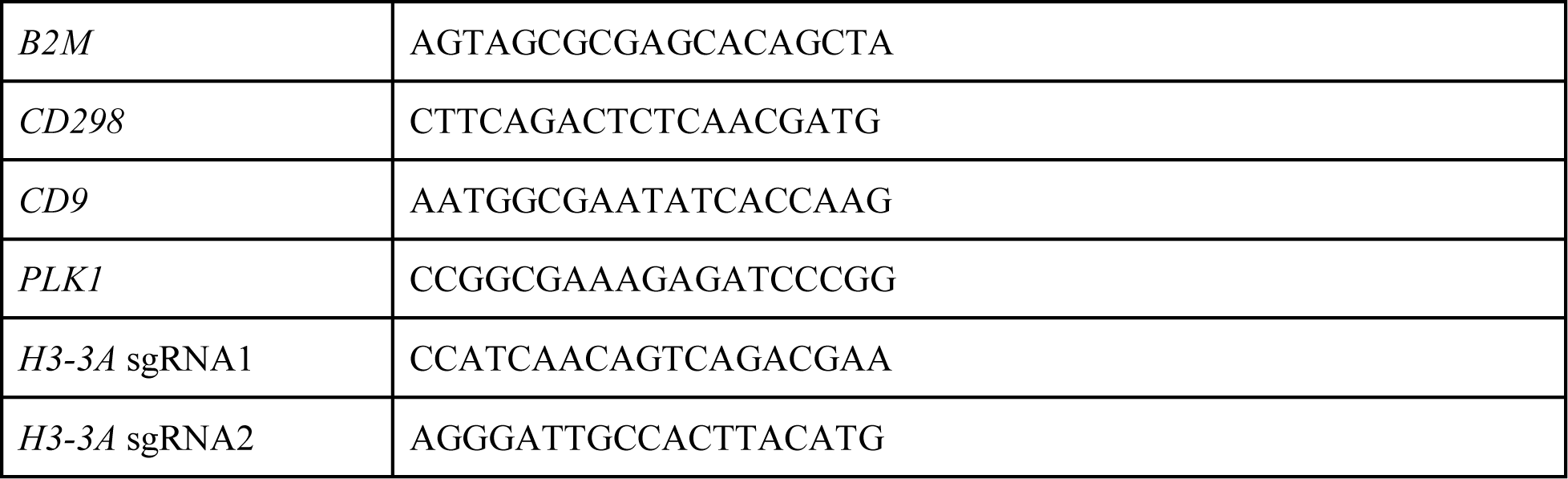
sgRNA sequences for CRISPRn.

**Table 3.**
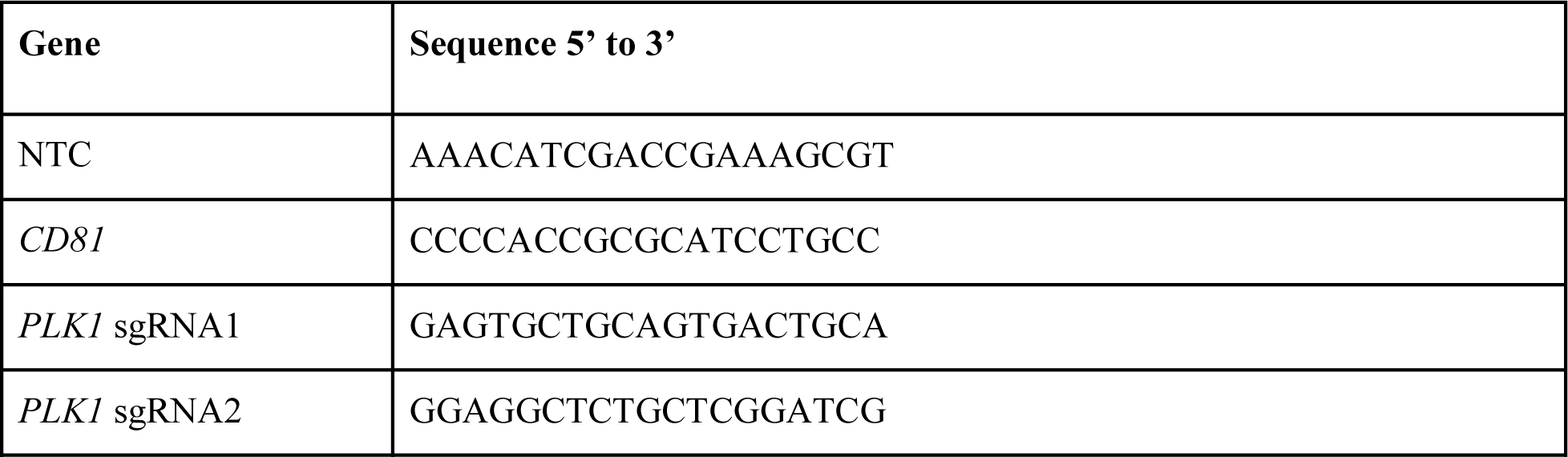
sgRNA sequences for CRISPRi.

**Table 4.**
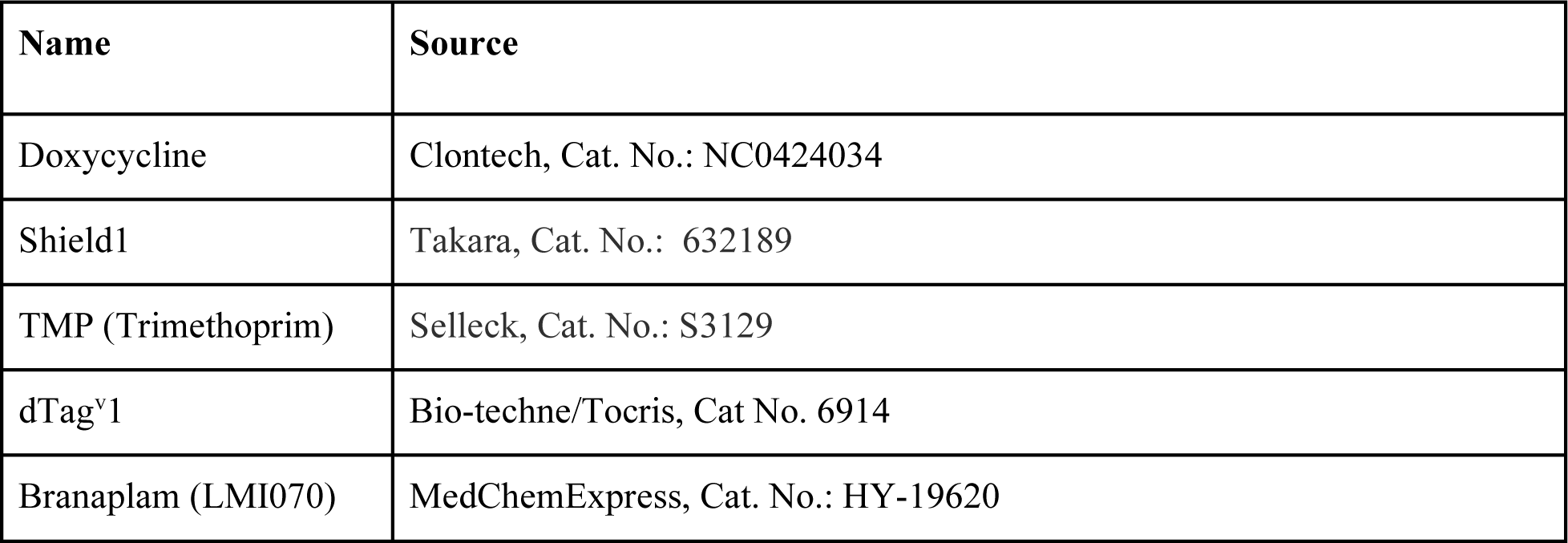
Small molecule drugs used in this study.

**Table 5:** Sequences of plasmids used in this study. <GSHEET attachment>

## Acknowledgements

We would like to thank Zhainib Amir for experimental assistance. We would also like to thank Soren Warming and Filip Roudnicky for helpful discussions and project support. Figures were prepared with BioRender.

This work was supported by Genentech, Inc., a member of the Roche group.

## Author Contributions

RS, TS, and BH conceptualized the project and designed experiments with input from AN. RS, TS, and BH designed reagents. RS, TS, AS, DW, LW, SX, AH, and CH performed the experiments. RS and TS analyzed the data and prepared figures. RS, TS, AN, and BH wrote the manuscript with input from all authors.

